# Genetic transformation and cell division delay in competent *Staphylococcus aureus*

**DOI:** 10.1101/2022.06.29.498089

**Authors:** Fedy Morgene, Célia Rizoug Zeghlache, Shi Yuan Feng, Yolande Hauck, Nicolas Mirouze

**Affiliations:** Université Paris-Saclay, CEA, CNRS, Institute for Integrative Biology of the Cell (I2BC), 9198, Gif-sur-Yvette, France

**Keywords:** *Staphylococcus aureus*, Horizontal gene transfer, Natural competence, Genetic transformation, cell cycle, division inhibition

## Abstract

Natural competence for genetic transformation, considered as one of the three main mechanisms leading to horizontal gene transfer in bacteria, is able to promote evolution, through genomic plasticity, and foster antibiotic resistance and virulence factors spreading. Conserved machineries and actors required to perform genetic transformation have been shown to accumulate at different cellular localizations depending on the model organism considered.

Here, we show in the human pathogen *Staphylococcus aureus* that DNA binding, uptake and recombination are spatially and temporally coordinated to ensure *S. aureus* genetic transformation. We also reveal that localization of genetic transformation proteins is dynamic and preferentially occurs in the vicinity of the division septum. We finally propose that *S. aureus* competent cells would initiate and then block cell division to ensure the success of genetic transformation before the final constriction of the cytokinetic ring.

## Introduction

Genetic transformation is one of the three main mechanisms allowing <u>H</u>orizontal <u>G</u>ene <u>T</u>ransfer (HGT) in prokaryotes. Genetic transformation refers to the ability displayed by certain bacteria to bind and take up exogenous DNA present in the environment for intracellular recombination with their own chromosome. This mode of HGT is clinically relevant as it is used by important human pathogens to acquire drug resistance or virulence genes (Griffith, 1928; Winter *et al.*, 2021). Interestingly, and at the opposite of conjugation and transduction, the two other main HGT mechanisms in bacteria, genetic transformation is entirely controlled by the recipient cell. Indeed, in order to express the proteins essential for genetic transformation, bacterial cells need to enter a specific physiological state called natural competence (Blokesch, 2016).

Many bacterial species have been shown to be able to enter the natural competence state to perform genetic transformation, one of the latest being the human pathogen *Staphylococcus aureus* (Morikawa *et al.,* 2012; Cordero *et al.,* 2022; Maree *et al.,* 2022). Under laboratory planktonic conditions, competence can be induced in late exponential phase in nearly 50% of *S. aureus* cells (Feng *et al.,* 2022). In addition, we have recently shown that competence is induced in response to a decrease in the oxygen concentration, an environmental stimulus that *S. aureus* often encounters during the course of an infection (Feng *et al.,* 2022). However, even though we have gained important insights in the competence regulatory network at play in *S. aureus*, almost nothing is known about the steps and actors defining genetic transformation in this human pathogen.

Importantly, the organization of the machineries accomplishing genetic transformation’s steps have been characterized in a number of model organisms (Dubnau and Blokesch, 2019). First, different structures have been shown to be involved in the initial exogenous DNA binding in various bacterial species. Indeed, while a competence-specific pilus, a structure evolutionary close from type IV pili and type-II secretion systems, might capture double-strand DNA (dsDNA) from the environment in *Streptococcus pneumoniae* (Laurenceau *et al.,* 2013), wall teichoic acids produced and modified during competence might represent the initial DNA binding site in *Bacillus subtilis* (Mirouze *et al.,* 2018). Uptake of dsDNA then occurs through binding to the membrane-anchored ComEA protein (Provvedi and Dubnau, 1999) but also requires the competence-specific pilus (Chung, Breidt and Dubnau, 1998), that might channel DNA through the cell wall to ComEA. The *comG* operon encodes most of the proteins required for the construction of this type IV transformation pilus (Chung and Dubnau, 1998). Assembly of the major pilin ComGC, and potentially additional minor pilins, all matured by the ComC peptidase (Chung and Dubnau, 1995), into this pilus requires the traffic ATPase ComGA (Haijema *et al.,* 2001). Once transported across the cell wall, exogenous dsDNA is converted to single-stranded DNA (ssDNA) (Lacks, Greenberg and Neuberger, 1975), which is then taken up into the cytosol through the aqueous transport channel formed by the ComEC integral membrane protein (Maier *et al.,* 2004; Draskovic and Dubnau, 2005). ComFA, a cytosolic ATPase containing helicase-like domains is thought to facilitate ssDNA internalization (Londono-Vallejo and Dubnau, 1994), through ComEC. Within the cell, the imported ssDNA is protected by a ssDNA-binding protein, SsbB, and the processing protein A, DprA (Berge *et al.,* 2003) which finally ensures the loading and polymerization of RecA (Quevillon-Cheruel *et al.,* 2012), to form presynaptic filaments leading to chromosomal recombinants.

Interestingly, localization of the transformation apparatus has shown that all the steps and actors involved are spatially and temporally closely organized (Hahn *et al.,* 2005; Kidane and Graumann, 2005). Indeed, exogenous dsDNA naturally binds in the direct vicinity of the proteins involved in its transport (Hahn *et al.,* 2005; Kidane and Graumann, 2005; Mirouze *et al.,* 2018). In addition, the proteins involved in DNA transport and recombination have also been shown to colocalize (Hahn *et al.,* 2005; Berge *et al.,* 2013). Importantly, all these steps and actors have been shown to be organized at different cellular locations in various model organisms: midcell in *S. pneumoniae* (Berge *et al.,* 2013) and at the cell pole in *B. subtilis* (Hahn *et al.,* 2005). Finally, despite those differences, the establishment of the transformation apparatus has often been associated to the cell cycle and the inhibition of cell division (Briley Jr., Prepiak, *et al.,* 2011; Bergé *et al.,* 2017).

In this study, we investigated the transformation apparatus composition, localization and dynamic in *S. aureus*. In particular, we show how ComGA localization is dynamic in space and time, evolving from diffuse in the cytosol, to associated to the inner face of the membrane, to finally concentrate in foci next to the division septum. In addition, ComGA localization seems coordinated with the cell cycle as most ComGA-expressing competent cells present a forming or completed septum, at the opposite of most non-competent cells. Importantly, ComGA localization evolves with the cell cycle, ultimately accumulating into foci in the direct vicinity of the septum. We finally confirm that representatives of each genetic transformation step co-localize with ComGA. All in all, we propose that the dynamic colocalization of genetic transformation machineries next to the septum have multiple roles: (i) to spatially coordinate DNA binding, uptake and processing, (ii) to synchronize these steps with *S. aureus* cell cycle and (iii) to impose a delay in cell division, probably through the inhibition of septum synthesis and/or constriction.

## Results

### ComGA exhibits several localization patterns in competent *S. aureus* cells

The traffic ATPase ComGA has been shown to be essential for both DNA binding and transport across the cell wall in *B. subtilis* (Briley Jr., Dorsey-Oresto, *et al.,* 2011). In addition, as it co-localizes with most of genetic transformation proteins, it has been used as a reference to visualize the transformation apparatus in numerous microscopy studies (Hahn *et al.,* 2005; Mirouze *et al.,* 2018). Therefore, we first investigated ComGA localization in *S. aureus* competent cells produced using our optimized “dilution” protocol (Feng *et al.,* 2022). To do so, we constructed a translational fusion between the gene encoding the green fluorescent protein (*egfp*) to the C-terminus of *comGA*, expressed from the natural *comG* promoter (St113, pRIT-P_*comG*_-comGA-egfp).

First, and as expected, ComGA-EGFP fluorescent signal was present in 19 to 23% of the total cells at the entry in stationary phase (**Supp. Fig. 1a**). Interestingly, in late exponential growth, ComGA-EGFP exhibited several localization patterns in distinct cells (**Fig. 1a**). Indeed, while in some cells ComGA-EGFP would be completely cytosolic, it was also found homogeneously localized or concentrated in individual foci at the inner face of the membrane in other competent cells (**Fig. 1b, Supp. Fig. 2 and Supp. Videos 1, 2 and 3**). Interestingly, in *B. subtilis*, ComGA has been shown to form individual or multiple foci associated to the inner face of the membrane (Hahn *et al.,* 2005). In comparison, it seemed that competent *S. aureus* cells would only form a single focus as cells displaying two or more foci only represented 0,8% of all the competent cells (**Supp. Table 1**).

**Figure 1.**
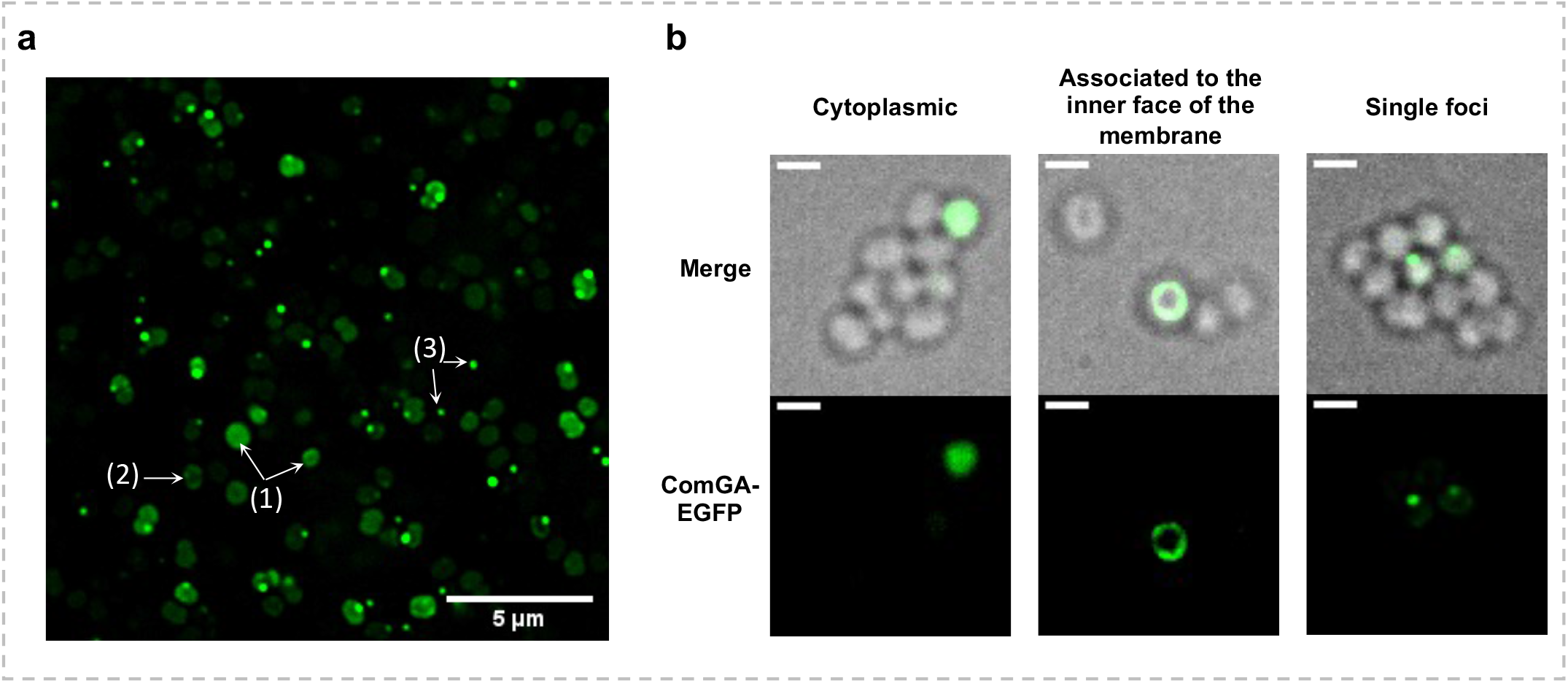
Localization of ComGA-EGFP in competent *S. aureus* cells. **(a)** St113 strain (pRIT-*P_comGA_-comGA;-egfp*) was grown to competence in CS2 medium (22h, 10^−5^ dilution). ComGA-EGFP displays several cellular localizations in *S. aureus* competent cells: cytoplasmic (1), associated to the inner face of the membrane (2) and accumulation in foci (3). Bar = 5 μm. **(b)** Examples of the three ComGA-EGFP cellular localizations observed in St113 cultures (pRIT-P*_comGA_-comGA-egfp*). 360 degrees and 3D rotations of these cellular localizations are presented in Supplementary Fig. 1 and Supplementary videos 1, 2 and 3. Bar = 1 μm.

### ComGA localization is dynamic

As we anticipated that ComGA needed to concentrate into foci to participate to the construction of the genetic transformation apparatus (Hahn *et al.,* 2005), we then wondered if ComGA localization could be temporally and spatially dynamic. To test this hypothesis, we analysed, along growth, the evolution of the different localization patterns of ComGA-EGFP (**Fig. 2**). Indeed, when competence started to develop (after 19h of growth), ComGA-EGFP was exclusively found diffuse in the cytoplasm of all the competent cells (**Fig. 2**). One hour later, cells where ComGA-EGFP localized at the inner face of the membrane started to appear. The following hour, ComGA-EGFP could be also observed as foci. Importantly, as the percentage of cells with foci increased, the percentage of cells with cytosolic or associated to the membrane localizations decreased accordingly (**Fig. 2**). Indeed, cells displaying a ComGA-EGFP focus represented nearly 60 % of the total competent cells after 25h of growth. This result was true for the three diluted cultures tested in which the final percentage of competent cells with a ComGA-EGFP focus was comprised between 57 and 75% (**Supp. Fig. 1b**).

**Figure 2.**
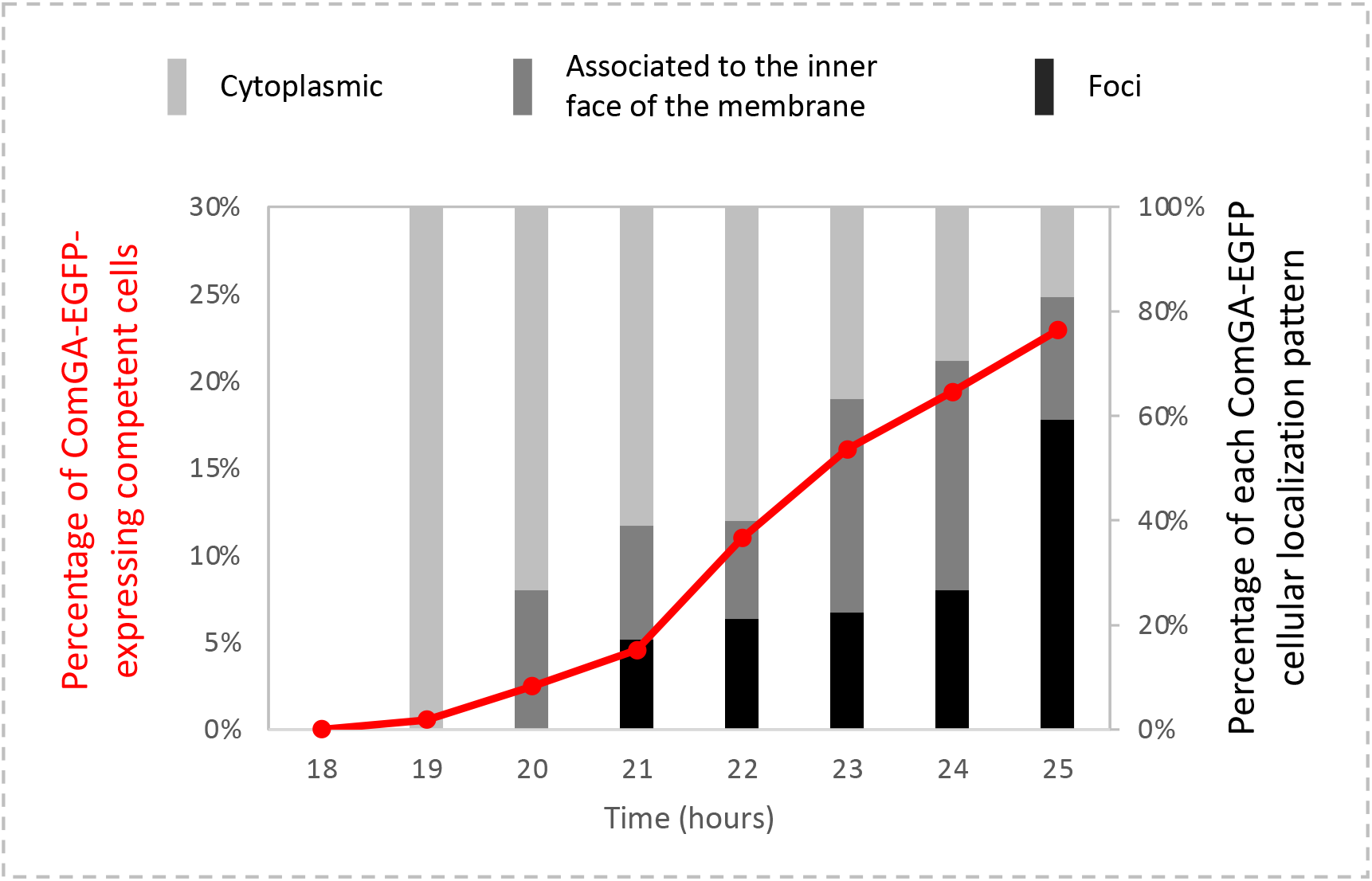
Spatial and temporal localization of ComGA is dynamic in competent *S. aureus* cells. Evolution of ComGA-EGFP localization patterns throughout growth was visualized using the St113 strain (pRIT-P*_comGA_-comGA-egfp*) grown in CS2 medium (10^−5^ dilution). Histograms represent the percentage of each ComGA-EGFP localization pattern between 18 and 25h of growth (cytoplasmic, light grey; associated to the inner face of the membrane, dark grey; single foci, black). The red curve represents the evolution of the percentage of ComGA-EGFP-expressing competent cells. At least 1500 cells were counted for each time point.

### ComGA foci preferentially localize next to the division septum

It was previously shown that the genetic transformation apparatus would localize at different cellular locations depending on the model organism considered (Hahn *et al.,* 2005; Berge *et al.,* 2013). Therefore, we then wondered if we could define a preferential cellular localization for the construction of the genetic transformation apparatus in *S. aureus* competent cells. To reach such goal, we constructed a *comGA-mcherry* translational fusion, expressed from the *comG* promoter and verified that it colocalized with the ComGA-EGFP fusion (**Supp. Fig. 3**). We then analysed the localization of ComGA-mCherry, in cells for which the cell wall was stained with Vancomycin BODIPY FL (Vanco-BODIPY, **Fig. 3**). In this experiment, the three ComGA localization patterns, and their dynamic, could be observed when fused to mCherry (**Supp. Fig. 4**). In addition, the Vanco-BODIPY staining allowed us to follow *S. aureus* cell division steps. Interestingly, recent observations have shown the morphological changes induced during *S. aureus* cell cycle (Monteiro *et al.,* 2015). These findings allowed us to correlate ComGA dynamic localization to the different stages of *S. aureus* cell division (**Fig. 3**).

**Figure 3.**
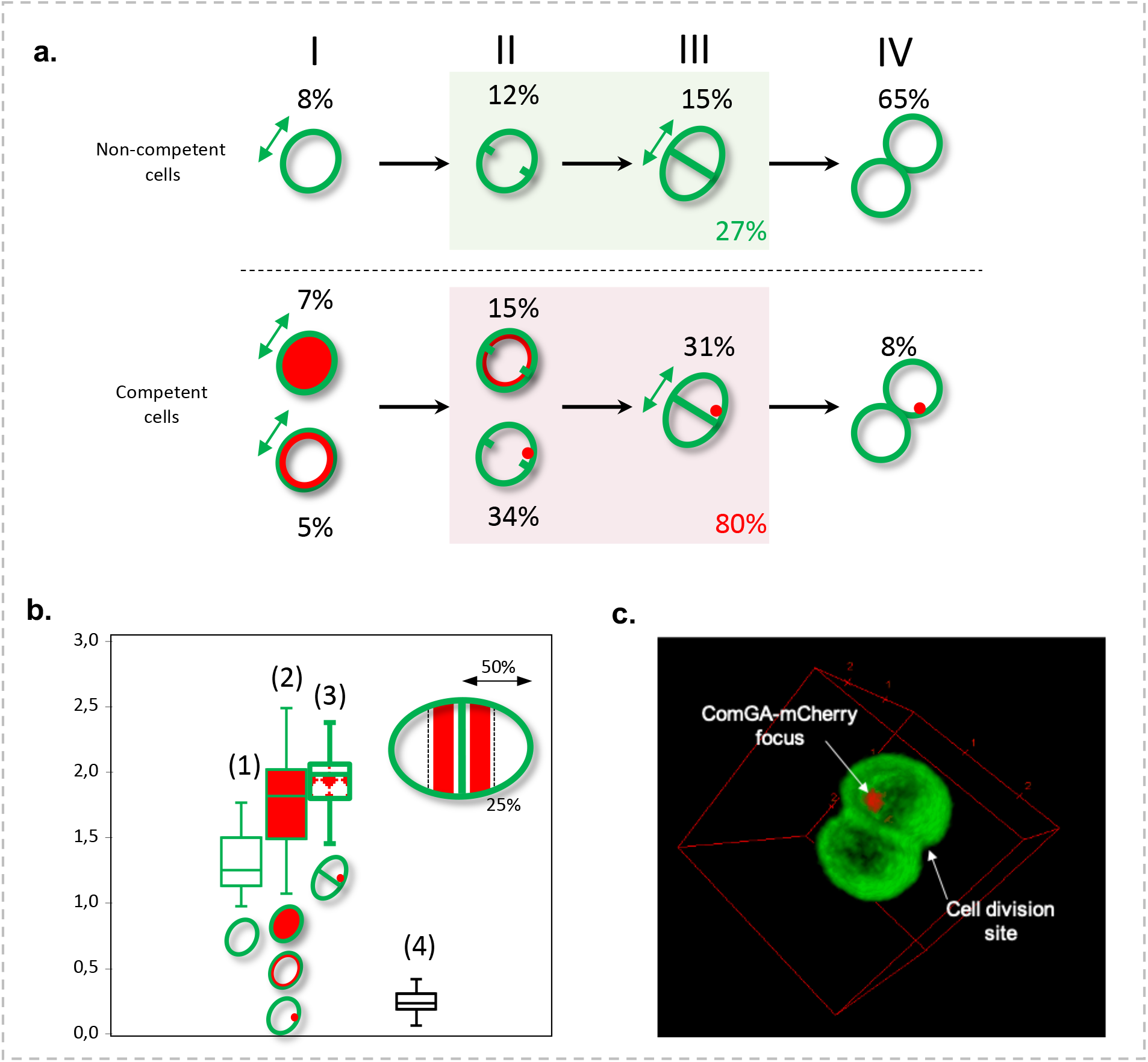
ComGA-mCherry forms foci near the division septum of *S. aureus* competent cells. **(a)** St228 (pRIT-P*_comGA_-comGA-mCh*) was grown for 25h (10^−5^ dilution) in CS2 medium and stained with Bodipy FL Vancomycin. Are presented the percentages of non-competent and competent cells in each step of cell division. In step I, single cells start elongating. Step II and III respectively correspond to septum initiation and completion. Note that cell elongation is promoted in step III. Finally, daughter cells splitting occurs in step IV. Figure adapted from (Monteiro *et al,* 2015) **(b)** Distribution of the long cell axis length for non competent (1) and competent (2 and 3) cells compared to the distance between ComGA foci and the septum (4). In (1), all the non-competent cells (dividing or not) have been considered. In (2) all the competent cells have been measured (dividing or not, with all the ComGA localization patterns). In (3), Only, the competent cells with ComGA-mCherry foci were considered. In (4) is presented the distribution of the distance between ComGA foci and the septum of the corresponding dividing competent cell. Distances were measured using the ImageJ software. In the top right corner of the graph is presented a drawing of a dividing competent cell, with the septum materialized at midcell. The red areas show the space where ComGA foci preferentially appear. **(c)** 3D reconstitution of a competent *S. aureus* cell presenting a single ComGA-mCherry focus localizing at the inner face of the membrane near the cell division site. (for 360 degrees rotation of this image, see supplementary video n°4).

First, we evaluated the percentage of non-competent and competent cells that were undergoing division (through the presence of a division septum, steps II and III in **Fig. 3a**). Under these conditions, only 27 % of the non-competent cells were found with a division septum (**Fig. 3a**). This relatively low number was expected as the culture approaches stationary phase and the generation time increases. At the opposite, 80% of the competent cells (i.e. expressing ComGA-Mcherry) were found with an initiated or completed division septum (**Fig. 3a**). Interestingly, this result was confirmed by the measurement of the non-competent and competent cells average size. Indeed, the average length of the long axis of all the non-competent cells (dividing or not) was evaluated at 1,32 ± 0,37 μm (**Fig. 3b**). This average length is in accordance with previous measurements of *S. aureus* cells (Monteiro *et al.,* 2015). In comparison, the average length of the long axis of all the competent cells reached 1,73 ± 0,36 μm (**Fig. 3b**). This result not only confirms that more competent cells are actively dividing but also indicates that dividing competent cells grow longer than dividing non-competent cells, potentially reflecting an arrest in competent cells division and an extended period of cell elongation (see step III in **Fig. 3a**).

Moreover, we showed that among the competent cells, cells that displayed ComGA-MCherry foci were the longest with an average length of the long axis reaching 1,93 ± 0,22 μm (**Fig. 3b**). This result allowed us to propose a hypothesis where ComGA dynamic localization could be coordinated with the steps of cell division. Indeed, we noticed that each ComGA localization pattern seemed to predominantly appear at different steps of division (**Fig 3a**). First, cells where ComGA-MCherry was cytoplasmic were mostly single cells (**Fig 3a**). Then, cells where ComGA-MCherry was uniformly associated to the inner face of the membrane were more often single cells or cells that just initiated division (**Fig 3a**). Following this dynamic, ComGA-Mcherry foci were found in dividing cells (where the division septum has been initiated or completed) and cells that just separated (**Fig 3a**). Therefore, we hypothesized that localization of ComGA foci, materializing the place where genetic transformation occurs, might be linked to the cell cycle.

To further investigate this last hypothesis, we finally calculated the distance between ComGA-MCherry foci and the division septum in competent cell. Interestingly, ComGA-MCherry foci were found, in average, 220 ± 90 nm away from the septum (**Fig. 3b, 3c, Supp. Fig. 4b and Supp. Video 4**). When compared to the average length of the long axis of competent cells with ComGA-MCherry foci (i.e. 1,93 μm), this result implies that the transformation apparatus is always localized in the direct vicinity of the division septum. Based on all these results, we provide in the discussion a model in which, during competence, cell division and genetic transformation are co-regulated in space and time to ensure the completion of a horizontal gene transfer event.

### Essential genetic transformation proteins co-localize with ComGA

Then, we decided to verify the localization of other important actors involved in genetic transformation, that were observed to co-localize with ComGA in historical model organisms (Hahn *et al.,* 2005). As the competence pilus has been shown to be essential for DNA binding and transport during genetic transformation (Chung and Dubnau, 1998; Laurenceau *et al.,* 2013), we first investigated its localization in competent *S. aureus* cells (**Fig. 4**). To do so, we inserted a FLAG tag at the C-terminus of ComGC, the major pilin. Importantly, three main types of localization (or structure) were observed (**Fig. 4a**). First, ComGC-FLAG was observed entirely associated to the membrane. This was not surprising, since ComGC is first anchored in the membrane and then liberated outside upon cleavage by ComC (Chung and Dubnau, 1995). In addition, in some cells ComGC could be observed as a single focus associated to the membrane/cell wall or as a single appendage, anchored in the membrane and extending in the environment (**Fig. 4a**). Interestingly, the staphylococcal transformation pilus displayed an average length of 0,9 ± 0,4 μm, a result comparable to what has been described in *S. pneumoniae* (Laurenceau *et al.*, 2013). Unfortunately, the competence-induced appendage length is usually difficult to assess because they often break during sample preparation (Laurenceau *et al.,* 2013). As a result, very few *S. aureus* competent cells with an intact transformation pilus could be observed (less than 1% of the cells with ComGC-FLAG signal).

**Figure 4.**
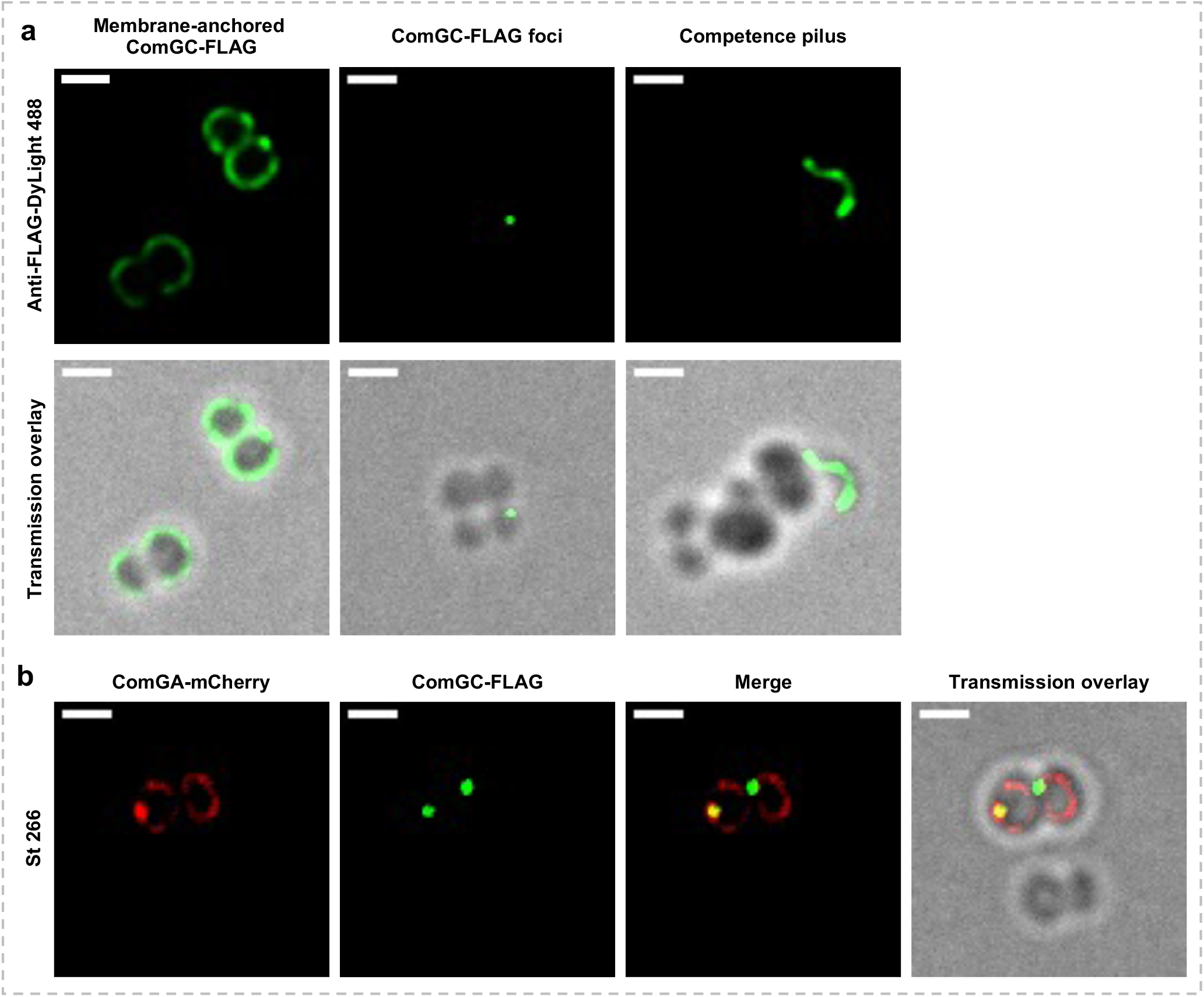
Localization of ComGC and the formation of the competence pilus in *S. aureus*. **(a)** ComGC-FLAG localization patterns in strain St243 (pRIT-P*_comGA_-comGC-FTAG*) grown in CS2 medium for 25h (10^−5^ dilution). From left to right: examples of membrane-anchored ComGC-FLAG, foci of ComGC associated to the membrane and ComGC-FLAG forming a pilus with an average size of 0,9 ± 0,4 μm. Bar = 1 μm **(b)** Colocalization of ComGA-mCherry and ComGC-FLAG in an overnight culture of strain St266 (pCNi-P*_comGA_-comGA-mCherry* / pRIT-P*_comGA_-comGC-FTAG*) in CS2 medium for 25 hours (10^−5^ dilution). Bar = 1 μm

We then chose to investigate the localization of proteins involved in (i) DNA transport (DNA binding, membrane anchored ComEA protein, the plasmic membrane pore forming protein, ComEC and the helicase-like protein, ComFA) and (ii) DNA processing (the RecA loader, DprA, and the single strand DNA binding protein, SsbB). N- or C-terminal translational fusions of these proteins fused with EGFP were made and expressed from their natural promoter. As expected, the membrane-associated proteins, ComEA, ComEC and ComFA were all found localizing throughout the membrane with ComEA and ComFA also accumulating as foci near the membrane (**Fig. 5a** and **Supp. Table 2**). Furthermore, the cytosolic DNA processing proteins, DprA and SsbB, were both found diffuse in the cytosol with some accumulation as foci near the membrane (**Fig. 5a** and **Supp. Table 2**). Interestingly, all the proteins examined above tend to accumulate in foci at the membrane (with the exception of ComEC), probably to coordinate genetic transformation’s steps.

**Figure 5.**
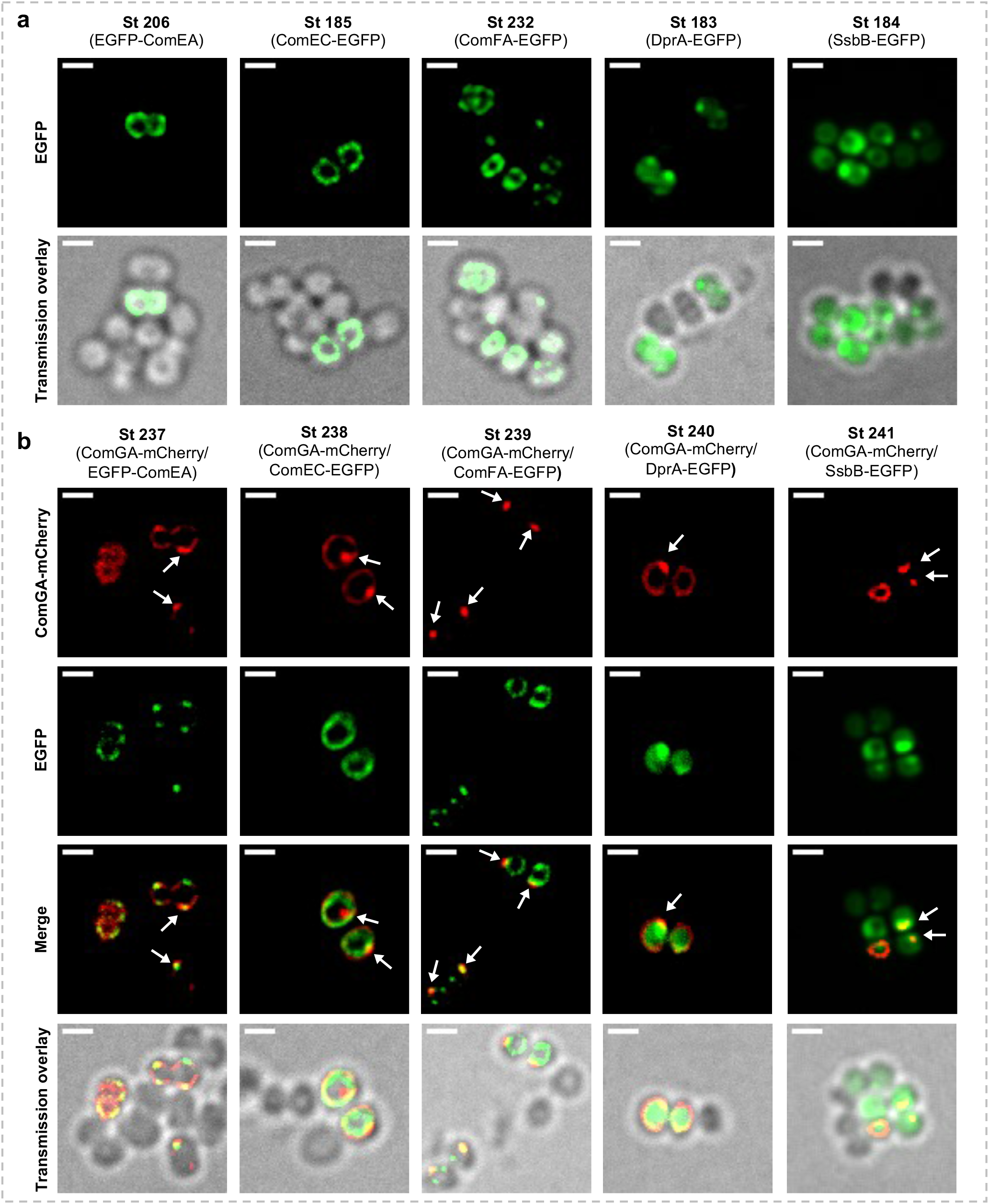
Co-localization of genetic transformation proteins in competent *S. aureus* cells. **(a)** Individual localization of ComEA (St206, pRIT-P*_comEA_-egfp-comEA*), ComEC (Stl85, pRIT-P_*comEc*-comEC-egfp_), ComFA (St232, pRIT-P*_comFA_-comEC-egfp*), DprA (Stl83, pRIT-P*_dprA_-dprA-egfp*) and SsbB (Stl84, pRIT-P*_ssbc_-ssbB-egfp*) all grown to competence for 25h in CS2 medium (10^−5^ dilution). Proteins involved in DNA binding and uptake (ComEA, EC and FA) all appeared associated to the membrane with ComEA and ComFA also accumulating in foci. The proteins involved in exogenous DNA processing (DprA and SsbB) were preferentially found in the cytoplasm but also accumulating in foci near the membrane. Bar = 1 μm. **(b)** ComGA colocalizes with other natural transformation apparatus proteins. Foci formed by ComEA (St237, pRIT-P*_comEA_-egfp-comEA* / pCNi-PcomGA-comGA-mCh), ComFA (St239, pRIT-P*_comFA_-comEC-egfp* / pCNi-PcomGA-comGA-mCh), DprA (St240, pRIT-P*_dprA_-dprA-egfp* / pCNi-PcomGA-comGA-mCh) and SsbB (St241, pRIT-P*_ssbc_-ssbB-egfp* / pCNi-PcomGA-comGA-mCh) co-localize with ComGA-Mcherry. Only ComEC (St238, pRIT-P*_comEc_-comEC-egfp* / pCNi-PcomGA-comGA-mCh) did not show any preferential co-localization with ComGA. Arrows indicate ComGA-mCherry foci. Bar : 1 μm.

In order to test this last hypothesis, we finally analysed the colocalization of all the proteins investigated above (i.e. ComGC, ComEA, ComEC, ComFA, DprA and SsbB) with that of ComGA-MCherry. First, we confirmed that ComGA is involved in the transformation pilus assembly as it was found co-expressed and colocalizing with the major pilin ComGC (**Fig. 4b** and **Supp. Table 3**). This result was predominantly obtained when ComGC formed foci (**Fig. 4b**). Similarly, each pair of transformation proteins fused to EGFP and ComGA-MCherry were found to be co-expressed in 83 to 98% of the cells, depending on the EGFP fusion considered (**Supp. Table 4**). Finally, we confirmed that all the transformation proteins fused to EGFP tested accumulated as foci near the membrane (with the exception of ComEC) at the exact same location as ComGA-Mcherry foci (**Fig. 5b**), probably reflecting the place where genetic transformation preferentially occurs.

### Exogenous DNA preferentially binds in the vicinity of the transformation apparatus

Finally, we wanted to visualize exogenous DNA binding at the surface of *S. aureus* competent cells. To do so, fluorescently-labelled (ATTO-550-dUTP) DNA was mixed with competent cells from different strains (**Fig. 6**). First, using a strain expressing *egfp* under the control of the *comG* promoter (as a reporter of competence development), we confirmed that DNA binding was highly specific to competent cells (**Fig. 6a** and **Supp. Table 5**). Indeed, 82± 7 % of the cells binding DNA were found to be competent (**Supp. Table 5**). Then, we particularly showed that fluorescently-labelled DNA preferentially binds in the vicinity of ComGA foci (**Fig. 6b** and **Supp. Table 5**). In addition, we evaluated that, in average, fluorescently-labelled DNA would localize 300 ± 100 nm from ComGA-EGFP. Finally, we observed that fluorescently-labelled DNA also preferentially binds to the genetic transformation pilus (**Fig. 6c** and **Supp. Table 5**). Again, due to the low number of cells harboring an intact pilus, this event was not often observed. However, this result is similar to what has been observed in *S. pneumoniae* (Laurenceau *et al.,* 2013) and could imply that in *S. aureus,* the transformation pilus also represents the preferential exogenous DNA binding site.

**Figure 6.**
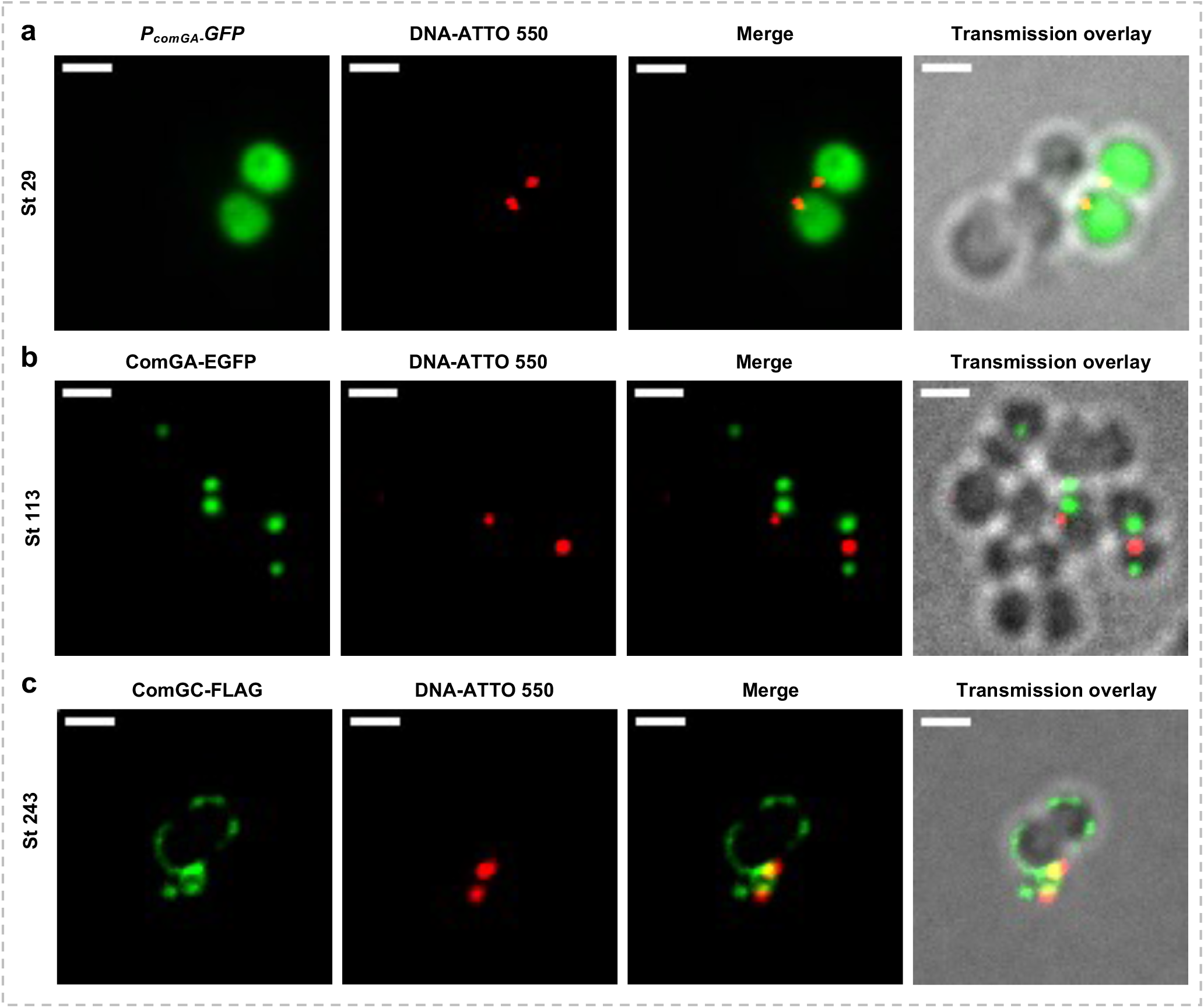
Exogenous DNA preferentially binds to *S. aureus* competent cells. **(a)** Exogenous fluorescently-labelled DNA (ATTO-550) preferentially binds to competent cells from the St29 strain expressing the *gfp* gene under the control of the *comG* promoter (pRIT-P*_comG_-gfp*) grown to competence in CS2 medium for 25h (10^−5^ dilution). Bar = 1 μm **(b)** Exogenous fluorescently-labelled DNA (ATTO-550) binds to St113 competent cells (pRIT-P*_comGA_-comGA-egfp*) near ComGA-EGFP foci grown in CS2 medium for 25h (10^−5^ dilution). Bar = 1 μm **(c)** Exogenous fluorescently-labelled DNA (ATTO-550) was found captured by the competence pseudopilus (containing ComGC-FLAG) stained with anti-FLAG-DyLight 488 in St243 cells (pRIT-P*_comGA_-comGC-FLAG*) grown to competence in CS2 medium for 25h (10^−5^ dilution). Bar = 1 μm

## Discussion

In *B. subtilis* competent cells, ComGA has been shown to first appear as diffuse in the cytoplasm and then to localize at the inner face of the membrane where it could be found nonuniformly throughout the cell with preferential accumulations at the pole(s) and/or at the septum (Hahn *et al.,* 2005). This spatial and temporal evolution of ComGA localization patterns clearly indicated that they are both dynamic and regulated. The polar accumulation of ComGA is competence-dependent and therefore candidates interacting with ComGA and directing it to specific cellular localizations must be the product of a late competence gene (Hahn *et al.,* 2005). Indeed, it has been proposed that ComEB, the second gene of the *comE* operon encoding a dCMP deaminase, could be the mediator for ComGA polar localization in *B. subtilis* (Burghard - Schrod, Altenburger and Graumann, 2020). Importantly, we observed the same kind of spatial and temporal dynamic in *S. aureus.* Therefore, it is also tempting to propose that ComGA localization in *S. aureus* could also be directed by a late competence protein. However, because ComGA displays a polar localization in *B. subtilis,* while it tends to accumulate next to the division site in *S. aureus,* ComGA localization mediator might be different in these two model organisms and further studies will be required to identify ComGA anchor in *S. aureus*.

Our observations also support the idea of the presence of multi-protein machineries, assembled at a unique cellular localization to perform binding, transport and processing of transforming DNA. The individual proteins we have studied here not only localized in similar patterns (with the exception of ComEC that did not form foci), but also showed colocalization when tagged with distinct fluorophores. ComGA (a traffic ATPase needed for dsDNA binding and required for the competence pilus assembly), colocalizes with ComGC (the major competence pilus’ pilin), ComEA (a dsDNA binding protein), ComFA (a protein required for DNA transport) and DprA or Ssb (two proteins involved in ssDNA processing). Importantly, while ComGC and ComEA are both exposed outside in the cell wall, ComGA and ComFA are predicted to be associated to the inner face of the membrane and DprA or Ssb supposed to be cytosolic. Therefore, proteins required at different steps during genetic transformation and targeted to different cellular compartments were found colocalized in competent cells. These observations suggest that binding, uptake and processing of transforming DNA are closely coordinated processes, in space and time, involving multiple protein-protein and DNA-protein interactions. Several reports have already showed that this was true in various model organisms and it will be essential to investigate this field in *S. aureus* in the future.

In addition, several genetic transformation proteins foci for an estimated 50 uptake sites per competent cell have been proposed in *B. subtilis*, suggesting that each of the fluorescent focus may contain several uptake machineries (Dubnau and Cirigliano, 1972). This might also be true in *S. aureus*.

However, the fact that almost all *S. aureus* competent cells displayed only one focus of transformation proteins might indicate that fewer uptake machineries might be assembled than in *B. subtilis*, potentially explaining, at least partially, the lower transformation efficiencies observed in this human pathogen (Feng *et al.*, 2022).

Very interestingly, our results also show that ComGA, and by extension the entire genetic transformation apparatus, preferentially localizes in the vicinity of *S. aureus* competent cells division septum (while most non-competent cells are not dividing). In addition, we establish a correlation between the dynamic of ComGA localization and the cell cycle stages. Therefore, we propose a model in which the genetic transformation apparatus’ localization, near the competent cell septum has a regulatory function in *S. aureus*: to establish a spatial and temporal link between DNA binding/uptake/processing and cell division.

In such model, competence-inducing *S. aureus* cells express ComGA which first appears diffuse in the cytoplasm (**step I in Fig. 3a**). As ComGA starts to uniformly associate with the inner face of the membrane, competent cells initiate division (**step II in Fig. 3a**). The initiation of the septum would then represent a spatial reference for ComGA accumulation, materializing the place where the transformation apparatus is established and genetic transformation occurs (**step II and III in Fig. 3a**). Finally, cell division completion generates two daughter cells with only one of them harboring a ComGA focus (**step IV in Fig. 3a**).

Finally, division inhibition during competence for genetic transformation has been demonstrated in several model organisms. For example, the early competence protein ComM delays division in the Gram-positive coccus, *S. pneumoniae* (Bergé *et al.,* 2017). The authors proposed that all these mechanisms could ensure genetic transformation completion before resumption of cell division.

Interestingly, it has been shown that during step III of division, *S. aureus* cells presenting a completed septum tend to elongate before septum constriction and daughter cells separation (Monteiro *et al.,* 2015). If septum synthesis and/or constriction is postponed in competent cells during step II and III (during which all the proteins co-localize to form the transformation apparatus), cell elongation could be prolonged, leading to longer competent cells and providing time for genetic transformation completion. Interestingly, initiation of constriction of the cytokinetic ring and completion of cell division have been shown to be delayed by ComM during genetic transformation in *S. pneumoniae* (Bergé *et al.,* 2017). It will important in the future to identify an early competence protein, similar to ComM from *S. pneumoniae,* regulating the cell cycle during genetic transformation in *S. aureus* competent cells.

## Acknowledgments

*S. aureus* N315 ex w/o φ (St12) and the staphylococcal natural competence reporter strain (St 29) were kindly donated by Pr Morikawa, Tsukuba University – Japan. Non-replicative plasmids pBCB-7-ChK and pBCB-8-GK, respectively encoding mCherry and EGFP, were a generous gift from Pr Pinho, NOVA University Lisbon – Portugal. High-copy-number *E. coli*-staphylococcal shuttle vector pCN34 was kindly obtained from Pr Novick, New York University Medical Center – USA.

This work was supported by a “Young Researcher grant” from the French National Research Agency to Nicolas Mirouze (ANR-18-CE35-0004 GenTranSa).

All microscopy acquisitions were performed at the Imagerie-GIF facility, I2BC in Gif-sur-Yvette – France.

## Author contributions

Conceptualization, F.M. and N.M.; Methodology, F.M. and N.M.; Investigation, F.M., C.R.Z., S.Y.F and Y.H.; Writing original draft, N.M.; Funding acquisition, N.M.; Resources, N.M.; Supervision, F.M. and N.M.

## Materials and Methods

### Bacterial strains and growth conditions

All bacterial strains and plasmids used in this study are listed in **Supp. Table 6**. *S. aureus* strains were grown at 37°C under aerobic conditions in Brain Heart Infusion (BHI) or Competence-inducing Synthetic medium (CS2). CS2 was freshly prepared from stock solutions as previously described (Morikawa *et al.,* 2012).

*E. coli* IM08B or DH10B strains were used as vector hosts to amplify plasmids and were cultured in Luria-Bertani (LB) medium at 37°C under aerobic conditions. When needed, one or several of the following antibiotics were added to the culture media depending on the bacterial genotype: 100 μg/mL ampicillin (Amp), 10 μg/mL chloramphenicol (Cm), 200 μg/mL kanamycin (Kan) and 50 μg/mL neomycin (Neo).

### Plasmids construction and transformations

Briefly, all the late competence genes studied in this work were amplified from the chromosomal DNA of *S. aureus* N315 ex w/o φ (St12) with their respective native promoters. **Supp. Table 7** presents more details about these genes and their promoters. Staphylococcal genomic DNA was extracted using the NucleoSpin® Microbial DNA extraction kit (Macherey-Nagel™) according to the producer’s protocol. Promoters, genes and linear plasmid vectors were amplified by PCR reaction using the Phusion™ High-Fidelity DNA Polymerase (Thermo Scientific™) and the appropriate primers following the manufacturer’s recommendations. The list of the primers is detailed in the **Supp. Table 8**. PCR products were checked by electrophoresis onto 1% agarose gel and purified by NucleoSpin® PCR and gel clean-up kit (Machery-Nagel™). Linear DNA fragments (inserts and vectors) with 3’ and 5’ overlapping regions were then assembled with a ratio of 1:3 (vector to insert) using the Gibson method with a homemade mixture containing 0.005 U/μL T5 Exonuclease, 0.03 U/μL Phusion® High-Fidelity DNA Polymerase and 5 U/μL T4 DNA Ligase in ISO buffer (5% w/v PEG-8000, 100 mM Tris.HCl, pH 7.5, 10 mM MgCl2, 10 mM DTT, 1 mM NAD and 1mM each dNTP). All reagents used for Gibson assembly were purchased from New England BioLabs™. The reaction was performed in a total volume of 20 μL for 1 hour at 50°C then inactivated by incubation for 10 minutes at 80°C. 5 μL of the Gibson assembly products were electroporated into 45 μL of electrocompetent *E. coli* IM08B (2.5 kV, 1 ms). Positive clones were selected on LB-Amp agar plates and checked by colony-PCR using DreamTaq Green PCR Master Mix (Thermo Scientific™) with the appropriate primers. Plasmids from selected positive clones were extracted and purified on silica columns (NucleoSpin® Plasmid, Macherey-Nagel™), then sequenced with their specific oligonucleotides (Eurofins Genomics). Finally, 500 ng of plasmids were used for transfer in electrocompetent *S. aureus* cells (1.8 kV, 2.5 ms). Transformed clones were selected on BHI agar plates with the appropriate antibiotics and checked by colony-PCR. Positive clones were cultured overnight and stored at −80°C in sterile cryogenic vials with 16% sterile glycerol.

### pRIT-P*_comG4_-cornGA-egfp*

It is important to mention that expression of the *comGA-egfp* translational fusion was preformed from the low copy number pRIT plasmid as no signal could be observed when expressed as a single copy on the chromosome. As *comGA* is one of the most induced gene during competence (Berka *et al.,* 2002; Fagerlund, Granum and Havarstein, 2014; Feng *et al.,* 2022), we also expressed the other translational fusions from the pRIT or pCBC low-copy number vectors.

In order to construct the plasmid pRIT-P*_comGA_-comGA-egfp*, a 1228 bp DNA fragment was first amplified from purified genomic DNA of N315 ex w/o φ (St 12) using the primers P152 and P220. This sequence contained the native promoter P*_comGA_* (256 bp) and the *comGA* gene without the stop codon (972 bp) flanked by KpnI restriction sites at both 5’ and 3’ termini. Purified KpnI-digested inserts were then ligated into KpnI-digested pBCB-8-GK plasmid and electroporated into *E. coli* IM08B. The pBCB-P*_comGA_-comGA-egfp* plasmid was purified, verified by PCR and sequenced to assure the correct orientation of the insert and the absence of mutations.

The P*_comGA_-comGA-egfp* DNA fragment was amplified from the pBCB-P*_comGA_-comGA-egfp* using primers P320 and P321 and assembled using the Gibson’s technique with the pRIT vector amplified using the primers P322 and P323. The resulting Gibson assembly was introduced into electrocompetent IM08B cells, and positive clones were selected on LB-Amp agar plates and checked by colony-PCR using the primers P335 and P336. The plasmid was finally extracted, purified and sequenced before electroporation into St 12 electrocompetent cells, thus creating the strain St113.

### Other pRIT-P_com_-com-egfp

pRIT-P*_comGA_-comGA-egfp* is a low-copy-number *E. coli-S. aureus* shuttle plasmid that allows the expression of ComGA under the control of its native promoter with a C-terminal EGFP translational fusion, mediated by a flexible 13 amino-acids linker. This plasmid served as an original template for the construction of all other pRIT plasmids used in this study using the same methodology except for the plasmid pRIT-P*_comEA_-egfp-comEA* in which the *egfp* translational fusion was designed at the N-terminal extremity of the *comEA* gene, separated by a 10 amino-acids linker.

### pRIT-P_comGA_-comGA-rnCh

With the aim of colocalizing ComGA-mCherry with other EGFP-fused late competence genes, we first designed the high-copy-number *S. aureus* replicative plasmid pCNi by replacing the erythromycin resistance cassette *ermC* of the plasmid pCN35 with the kanamycin resistance cassette *aphA-3* from the plasmid pCN34 using ApaI and SacII restriction enzymes (FastDigest™, Thermo Scientific®). The resulting pCNi plasmid contained the staphylococcal origin of replication pT181*cop*-623 *repC* and the kanamycin resistance cassette *aphA-3.* pCNi was first produced in DH10B cells then transferred into RN 4220 and finally into St12 thus creating the strain St224. Using the Gibson assembly technique, we created the plasmid pRIT-P*_comGA_-comGA-mCh* by replacing the *egfp* gene of th

### Natural competence development

The desired *S. aureus* strains were inoculated onto BHI agar plates, with the appropriate antibiotics if needed, and incubated at 37°C for 24 hours. CS2 suspensions adjusted to OD_600 nm_ of 0.5 were prepared from liquid BHI pre-cultures in exponential phase of growth. Serial 10-fold dilutions were then prepared in a total volume of 10 mL in individual sterile 50 mL Falcon tube and incubated overnight at 37°C with shaking at 120 rpm. Cell density was monitored hourly and 500 μL samples were collected in sterile Eppendorf tubes. Cell pellets were washed with sterile PBS, fixed with 1.6% formaldehyde (Invitrogen™) for 20 minutes at room temperature, rinsed twice in equal volume of sterile PBS and finally re-suspended in 100 μL of 45% glycerol buffer for storage at −20°C.

### Spinning disk microscopy

10 μL of fixed cells were mounted on 1% agarose coated slide with a drop of mounting medium (Fluoromount-G®). Slides were then observed under a confocal microscope Nikon Eclipse Ti-E equipped with a spinning disk module (Yokogawa CSU-X1-A1) and a super resolution system (Live SR 3D, Gataka systems) equipped with a x100 oil immersion objective (Nikon; NA = 1.49). Fluorescence images were recorded using the 470/30-nm excitation filter and 520/35-nm filter for GFP, the 560/40-nm excitation filter and 630/60-nm filter for mCherry. Images were processed with The Metamorph software (Molecular Devices). Images were finally analyzed with the FiJi software (ImageJ 1.53c).

### Vancomycin, BODIPY™ FL Conjugate (VMB) labeling

Cells were labeled with Vancomycin BODIPY FL (Vanco-BODIPY, Life Technologies) at a final concentration of 2 μg/ml for 30 min at 37°C. Excess of incorporated labeling was removed by washing the cells three times in sterile PBS and finally fixed as explained above.

### Competence pseudopilus visualization

In order to observe the staphylococcal competence pseudopilus, 1 mL of overnight cultures in CS2 of strains expressing ComGC-FLAG was collected in a sterile Eppendorf tube, centrifuged at 4000 g for 3 minutes and washed with sterile PBS after. Cells were fixed directly on SuperFrost Plus™ slides with 3.2% PFA and labeled with anti-FLAG-DyLight 488 (1:200, ThermoFisher).

### Fluorescent DNA preparation

For DNA binding experiments, a 1kb fluorescent DNA was first synthesized by PCR using Aminoallyl-dUTP-ATTO-550 (JenaBioscience). Briefly, we used 1 μL of Extaq enzyme from Takara on genomic DNA (20 μg) with 1 μL each primer, 0,25 μM of each dNTP and 0,1 μM of ATTO-550 dUTP. Elongation was performed at 72°C for 3min per kb, fragments were purified (Machery-Nagel™) and assessed by spectrophotometry to check the incorporation of fluorescent nucleotides.

250 μL of overnight *S. aureus* cultures in CS2 were collected into a sterile tube, washed, resuspended in an equal volume of filtered CS2, preheated to 37 °C, and containing 500 ng of fluorescent DNA and incubated for 5 minutes at 37°C with gentle shaking. Samples were finally fixed as mentioned above.

