## Supplemental data for "Genetic transformation and cell division delay in competent *Staphylococcus aureus*"

**This PDF file includes:**

Supplementary figures 1 to 4
Supplementary tables 1 to 8
Supplementary videos 1 to 4

### Supplementary Figures

**Supplementary Fig. 1. Spatial and temporal dynamic of ComGA-EGFP localization in competent *S. aureus* cells.**

**(a)** Evolution of the percentage of ComGA-EGFP-expressing cells. Samples from serial 10-fold dilutions (10-3 in green, 10-4 in blue and 10-5 in red) of strain St113 (pRIT-PcomGA-comGA-egfp) grown in CS2 medium were collected every hour and analyzed microscopically for ComGA-EGFP expression.

**(b)** Dynamic of ComGA-EGFP localization patterns in *S. aureus* competent cells from diluted cultures (10-3 top panel; 10-4, middle panel; 10-5, bottom panel). In addition of the evolution of the percentage of ComGA-EGFP-expressing cells (10-3 in green, 10-4 in blue and 10-5 in red), histograms represent the percentage of ComGA-EGFP localizing in the cytoplasm (light grey), associated to the membrane (dark grey) or as single foci (black).

At least 1500 cells were counted for each time point in each culture.

**Supplementary Fig. 2. ComGA-EGFP localization patterns in competent *S. aureus* cells.**

Examples of the three ComGA-EGFP (pRIT-$P_{comGA}$-*comGA-egfp*) cellular localizations observed using the St113 strain grown in CS2 medium (from left to right: cytoplasmic, associated to the inner face of the membrane and accumulation in foci near the membrane).

For each localization pattern, we provide 3D reconstructions (top row) as well as 360 degrees videos.

Bar = 4 µm

**Supplementary Fig. 3. Colocalization of ComGA-EGFP and ComGA-mCherry in competent *S. aureus* cells.**

Samples from strain St242 (pRIT-$P_{comGA}$-*comGA-egfp* / pCNi-$P_{comGA}$-*comGA-mCherry*) grown in CS2 medium (dilution $10^{-3}$ for 24 hours) show co-localization of the two fusion proteins.

Bar = 2 µm

**Supplementary Fig. 4. Localization of ComGA-MCherry in *S. aureus* competent cells.**

**(a)** Examples of the three localizations of ComGA-Mcherry (cytoplasmic, top panel; associated to the inner face of the membrane, middle panel and single foci, bottom panel) in *S. aureus* competent cells (St228, *pCNi-$P_{comGA}$-comGA-mCh* ) stained with Bodipy FL Vancomycin.

Bar = 1 µm

**(b)** Examples of ComGA-MCherry (St228, *pCNi-$P_{comGA}$-comGA-mCh* ) foci localizing next to the initiated septum in *S. aureus* competent cells.

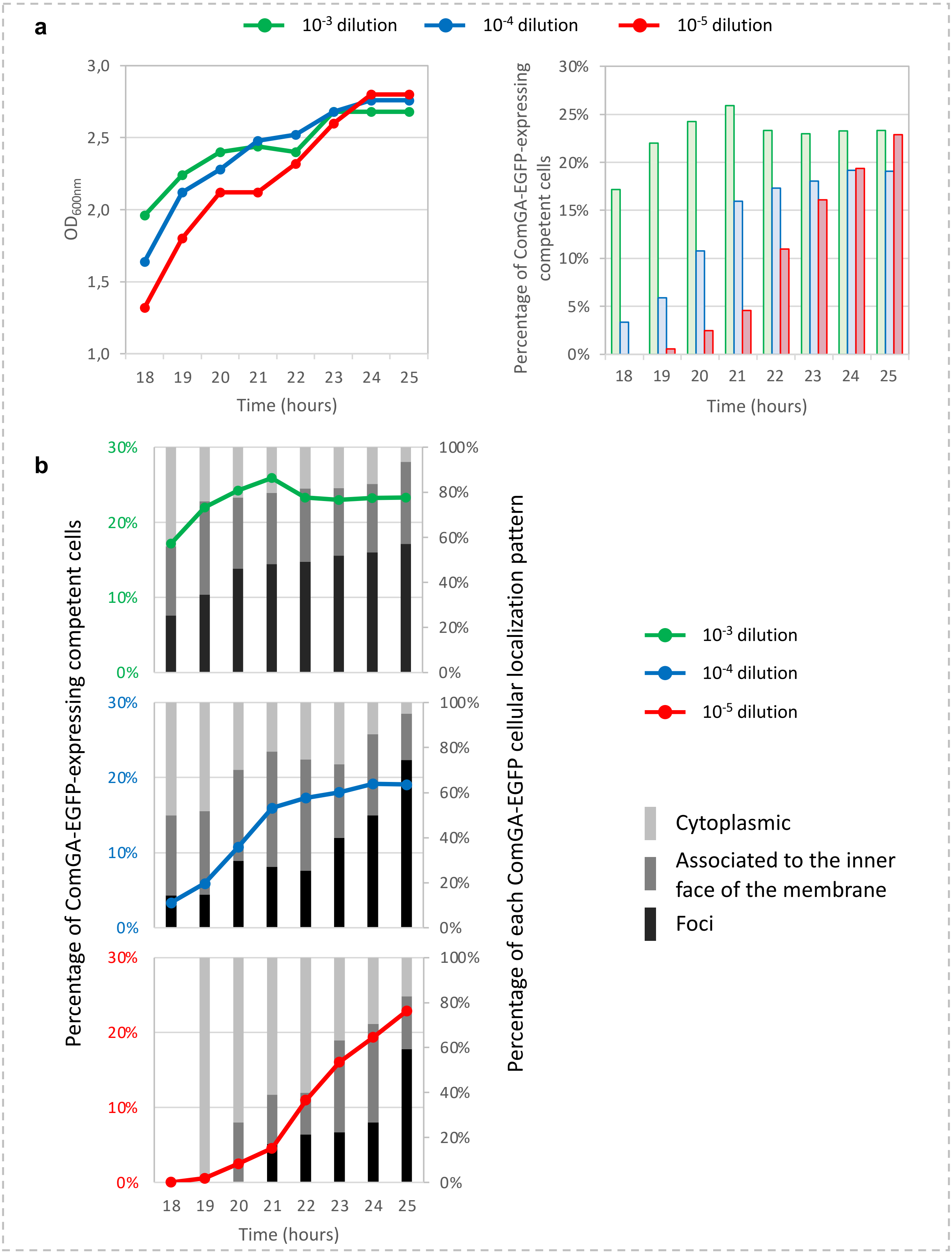


**Supplementary Fig. 1. Spatial and temporal dynamic of ComGA-EGFP localization in competent *S. aureus* cells.**

**(a)** Evolution of the percentage of ComGA-EGFP-expressing cells. Samples from serial 10-fold dilutions ($10^{-3}$ in green, $10^{-4}$ in blue and $10^{-5}$ in red) of strain St113 (pRIT-P$_{comGA}$-*comGA-egfp*) grown in CS2 medium (left panel) were collected every hour and analyzed microscopically for ComGA-EGFP expression (right panel).

**(b)** Dynamic of ComGA-EGFP localization patterns in *S. aureus* competent cells from diluted cultures ($10^{-3}$ top panel; $10^{-4}$, middle panel; $10^{-5}$, bottom panel). In addition to the evolution of the percentage of ComGA-EGFP-expressing cells ($10^{-3}$ in green, $10^{-4}$ in blue and $10^{-5}$ in red), histograms represent the percentage of each ComGA-EGFP localizing pattern : in the cytoplasm (light grey), associated to the membrane (dark grey) or as single foci (black). At least 1500 cells were counted for each time point in each culture.

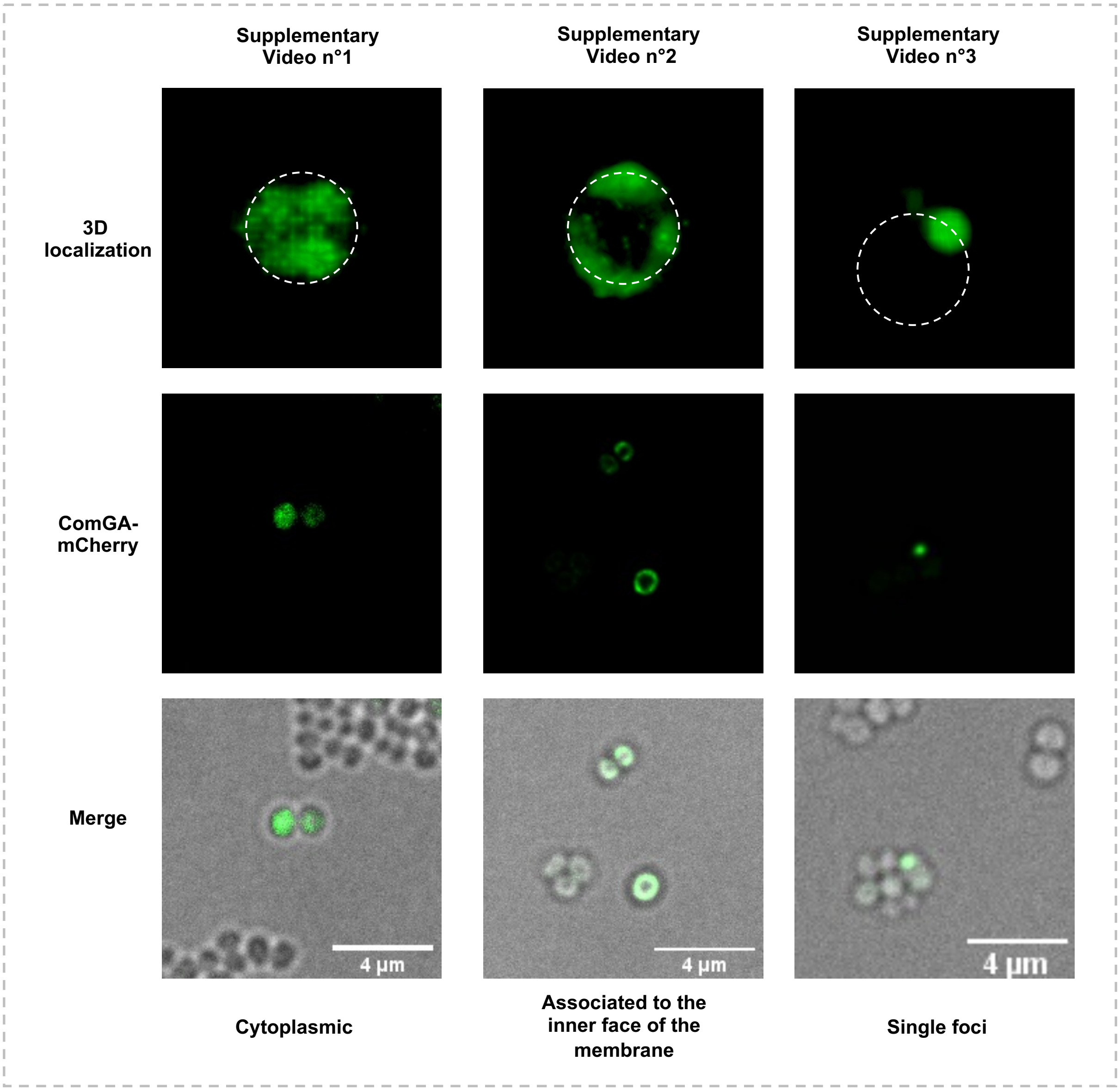


**Supplementary Fig. 2. ComGA-EGFP localization patterns in competent *S. aureus* cells.**
Examples of the three ComGA-EGFP (pRIT-$P_{comGA}$-*comGA-egfp*) cellular localizations observed using the St113 strain grown in CS2 medium (from left to right: cytoplasmic, associated to the inner face of the membrane and accumulation in foci near the membrane).
For each localization pattern, we provide 3D reconstructions (top row) as well as 360 degrees videos.
Bar = 4 µm

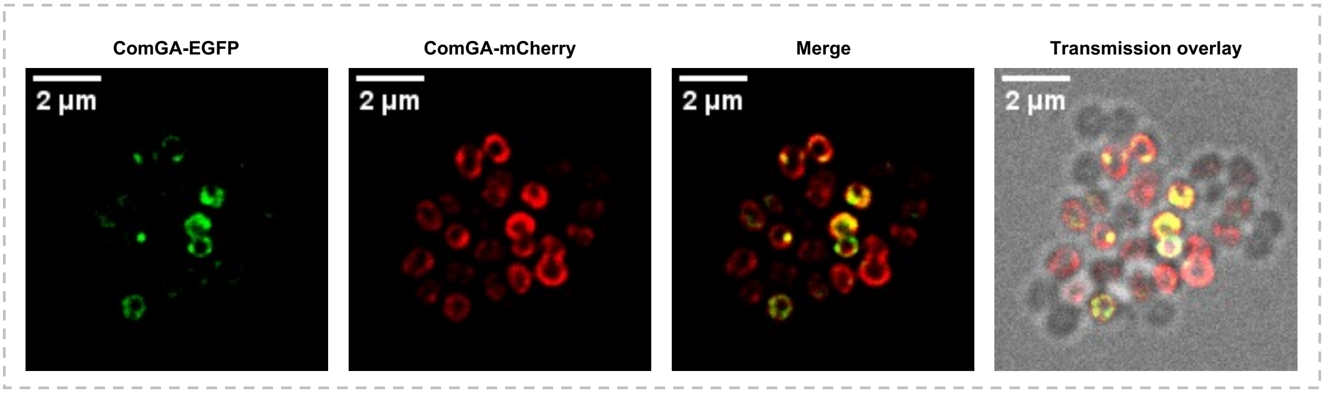


**Supplementary Fig. 3. Colocalization of ComGA-EGFP and ComGA-mCherry in competent *S. aureus* cells.**

Samples from strain St242 (pRIT-$P_{comGA}$-*comGA-egfp* / pCNi-$P_{comGA}$-*comGA-mCherry*) grown in CS2 medium (dilution $10^{-3}$ for 24 hours) show co-localization of the two fusion proteins.

Bar = 2 µm

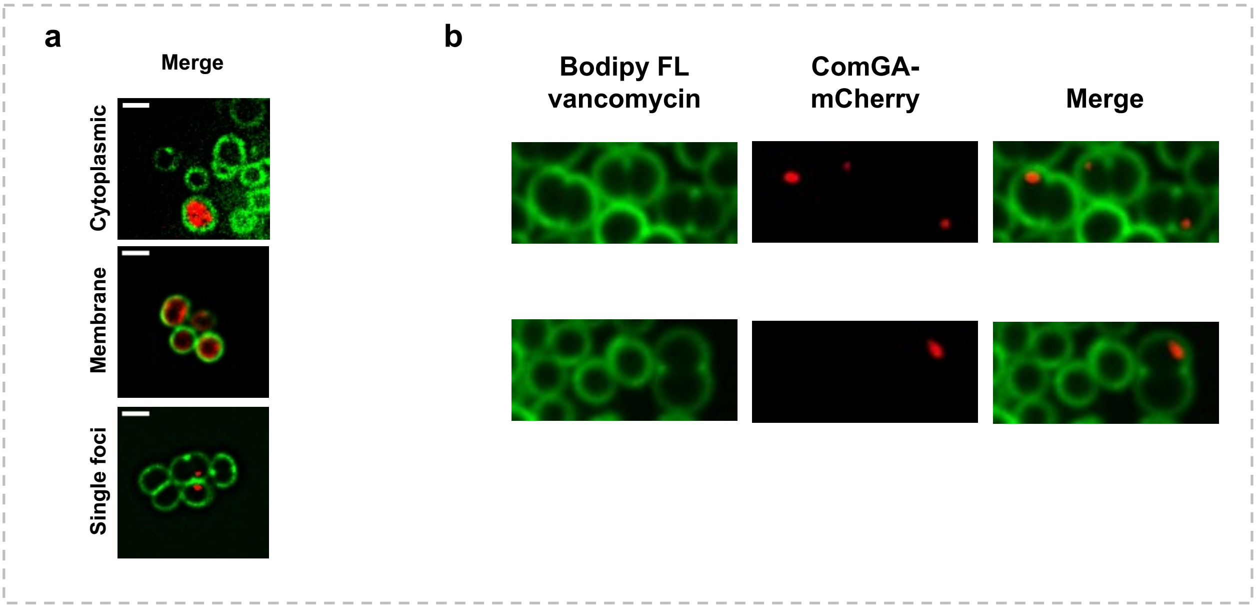


**Supplementary Fig. 4. Localization of ComGA-MCherry in *S. aureus* competent cells.**
**(a)** Examples of the three localizations of ComGA-Mcherry (cytoplasmic, top panel; associated to the inner face of the membrane, middle panel and single foci, bottom panel) in *S. aureus* competent cells (St228, $pCNi\text{-}P_{comGA}\text{-}comGA\text{-}mCh$ ) stained with Bodipy FL Vancomycin.
Bar = 1 µm
**(b)** Examples of ComGA-MCherry (St228, $pCNi\text{-}P_{comGA}\text{-}comGA\text{-}mCh$ ) foci localizing next to the initiated septum in *S. aureus* competent cells.

### Supplementary Tables

**Supplementary Table 1. ComGA-EGFP localization in *S. aureus* competent cells.**

| Total counted cells | % of cells with ComGA-EGFP | % of cells with ComGA foci | % of cells with a single ComGA focus | % of cells with two ComGA foci | % of cells with more than two ComGA foci |
|---|---|---|---|---|---|
| 13664 | 1548 (11.3%) | 478 (31.0%) | 474 (99.2%) | 3 (0.6%) | 1 (0.2%) |

Strain St113 (pRIT-P$_{comGA}$-*comGA-egfp*) was grown to competence for 24h in CS2 medium ($10^{-3}$ dilution). The percentage of ComGA-EGFP-expressing cells and among those the percentage of cells with ComGA-EGFP foci were evaluated by microscopy. Finally, among the cells presenting ComGA-EGFP foci, the percentage of cells with one, two or more foci was calculated.

This table completes the results presented in Fig. 1.

At least 400 cells were counted for each strain.

**Supplementary Table 2. Statistics for ComEA, ComEC, ComFA, DprA and SsbB localization in competent *S. aureus* cells.**

| Strain | St 206 | St 185 | St 232 | St 183 | St 184 |
|---|---|---|---|---|---|
| EGFP translational fusion | EGFP-ComEA | ComEC-EGFP | ComFA-EGFP | DprA-EGFP | SsbB-EGFP |
| Total EGFP+ % | 18% | 9% | 36% | 30% | 63% |
| Cytoplasmic | 0% | 0% | 0% | 69% | 45% |
| Membrane | 55% | 100% | 85% | 0% | 0% |
| Membrane foci | 45% | 0% | 15% | 31% | 55% |

Strains St206 (pRIT-$P_{comEA}$-*egfp-comEA*), St185 (pRIT-$P_{comEC}$-*comEC-egfp*), St232 (pRIT-$P_{comFA}$-*comEC-egfp*), St183 (pRIT-$P_{dprA}$-*dprA-egfp*) and St184 (pRIT-$P_{ssbC}$-*ssbB-egfp*) were grown to competence for 25h in CS2 medium ($10^{-5}$ dilution). The percentage of cells expressing each EGFP fusion as well as the percentage of each localization pattern were evaluated by microscopy.

This table completes the results presented in Fig. 5.

At least 400 cells were counted for each strain.

**Supplementary Table 3. ComGC-FLAG localization in *S. aureus* competent cells.**

| Strain | Counted cells | %ComGA | %ComGC | % ComGC+ in ComGA | % ComGA+ in ComGC |
|---|---|---|---|---|---|
| St 266 | 474 | 42% | 28% | 56% | 85% |

Strain St266 (pCNi-$P_{comGA}$-*comGA-mCherry* / pRIT-$P_{comGA}$-*comGC-FLAG*) was grown to competence for 25h in CS2 medium ($10^{-5}$ dilution). The percentage of ComGA-mCherry-expressing cells, ComGC-FLAG-expressing cells as well as the percentage of ComGA-mCherry-expressing cells that also express ComGC-FLAG and ComGC-FLAG-expressing cells that also express ComGA-mCherry were evaluated by microscopy.

This table completes the results presented in Fig. 4.

At least 400 cells were counted for each strain.

**Supplementary Table 4. ComEA, ComEC, ComFA, DprA and SsbB co-localization with ComGA-Mcherry in *S. aureus* competent cells.**

| **Strain** | **St 237** | **St 238** | **St 239** | **St 240** | **St 241** | **St 242** |
|---|---|---|---|---|---|---|
| EGFP translational fusion | EGFP-ComEA | ComEC-EGFP | ComFA-EGFP | DprA-EGFP | SsbB-EGFP | ComGA-EGFP |
| Total counted cells | 415 | 429 | 438 | 531 | 424 | 417 |
| % of ComGA-mCherry-expressing cells | 32% | 50% | 39% | 47% | 31% | 48% |
| % of EGFP-expressing cells | 17% | 41% | 33% | 34% | 60% | 34% |
| % of cells with EGFP fusions localization in ComGA-mCherry-expressing cells | 46% | 78% | 79% | 69% | **98%** | 69% |
| % of cells with ComGA-mCherry localization in EGFP-expressing cells | **83%** | **95%** | **92%** | **94%** | 51% | **98%** |

Strains St237 (pRIT-P$_{comEA}$-*egfp-comEA / pCNi-P$_{comGA}$-comGA-mCh*), St238 (pRIT-P$_{comEC}$-*comEC-egfp / pCNi-P$_{comGA}$-comGA-mCh*), St239 (pRIT-P$_{comFA}$-*comEC-egfp* / pCNi-PcomGA-comGA-mCh), St240 (pRIT-P$_{dprA}$-*dprA-egfp / pCNi-P$_{comGA}$-comGA-mCh*), St241 (pRIT-P$_{ssbC}$-*ssbB-egfp / pCNi-P$_{comGA}$-comGA-mCh*) and St242 (pRIT-P$_{comGA}$-*comGA-egfp / pCNi-P$_{comGA}$-comGA-mCh*) were grown to competence for 25h in CS2 medium ($10^{-5}$ dilution). The percentage of cells expressing ComGA-Mcherry, expressing each EGFP fusion as well as the percentage of ComGA-mCherry-expressing cells that also express the EFGP fusion and EGFP fusion-expressing cells that also express ComGA-mCherry were evaluated by microscopy.

This table completes the results presented in Fig. 5.

At least 400 cells were counted for each strain.

**Supplementary Table 5. Exogenous DNA binding at the surface of *S. aureus* competent cells.**

| Strain | Counted cells | %GFP | %DNA | %GFP+ in DNA+ | %DNA+ in GFP+ |
|---|---|---|---|---|---|
| St29 | 408 | 48% | 14% | **82%** | 23% |

| Strain | Counted cells | %ComGA-EGFP | %DNA | %GFP+ in DNA+ | %DNA+ in ComGA-EGFP+ |
|---|---|---|---|---|---|
| St113 | 401 | 31% | 12% | **80%** | 31% |

| Strain | Counted cells | %ComGC-FLAG+ | %DNA | %FLAG+ in DNA+ | %DNA+ in ComGC-FLAG+ |
|---|---|---|---|---|---|
| St243 | 427 | 23% | 10% | **88%** | 38% |

Strains St29 (pRIT-P$_{comGA}$-*egfp*), St113 (pRIT-P$_{comGA}$-*comGA-egfp*) and St243 (*pRIT-PcomGA-comGC-flag*) were grown to competence for 23h in CS2 medium ($10^{-5}$ dilution). After exposition to fluorescently labelled DNA, the percentage of competent cells, of cells binding DNA were evaluated. In addition, the percentage of cells binding DNA that were also competent as well as competent cells that were also binding DNA was calculated (two last columns).
This table completes the results presented in Fig. 6.
At least 400 cells were counted for each strain.

#### Supplementary table 6. Bacterial strains and plasmids

| Bacteria strains | | | | |
|---|---|---|---|---|
| **Strain** | **Genotype / plasmid** | **Phenotype** | **Parental** | **Source** |
| ***Escherichia coli*** | | | | |
| **IM08B** | Δ*dcm (*does not methylate DNA on cytosine*)* ΩP$_{help}$-*hsdMS* ΩP$_{N25}$-*hsdS* (type 1 modification gene from *S. aureus* CC8-2 and CC8-1, respectively) | Stp$^R$ | DC10B | (Monk *et al.*, 2015) |
| **DH10B** | F$^-$ Δ*mcr*A ϕ80*lac*ZΔM15 Δ*lac*X74 Δ*rec*A1 Δ*end*A1 *ara*D139 Δ(*ara leu*) λ$^-$ | Stp$^R$ | K-12 | (Grant *et al.*, 1990) |
| ***Staphylococcus aureus*** | | | | |
| **St 12** | N315 ex w/o ϕ (lacking *SCCmec* and cured from phages) | Erm$^R$ | N315 | (Morikawa *et al.*, 2012) |
| **St 29** | N315 ex w/o ϕ / pRIT-P$_{comGA}$-*gfp* | Erm$^R$, Cm$^R$ | St 12 | (Morikawa *et al.*, 2012) |
| **St 113** | N315 ex w/o ϕ / pRIT-P$_{comGA}$-*comGA-egfp* | Erm$^R$, Cm$^R$ | St 12 | This study |
| **St 183** | N315 ex w/o ϕ / pRIT-P$_{dprA}$-*dprA-egfp* | Erm$^R$, Cm$^R$ | St 12 | This study |
| **St 184** | N315 ex w/o ϕ / pRIT-P$_{ssbC}$-*ssbB-egfp* | Erm$^R$, Cm$^R$ | St 12 | This study |
| **St 185** | N315 ex w/o ϕ / pRIT-P$_{comEC}$-*comEC-egfp* | Erm$^R$, Cm$^R$ | St 12 | This study |
| **St 206** | N315 ex w/o ϕ / pRIT-P$_{comEA}$-*egfp-comEA* | Erm$^R$, Cm$^R$ | St 12 | This study |
| **St 228** | N315 ex w/o ϕ / pCNi-P$_{comGA}$-*comGA-mCh* | Erm$^R$, Kan$^R$ | St 12 | This study |
| **St 232** | N315 ex w/o ϕ / pRIT-P$_{comFA}$2-*comFA-egfp* | Erm$^R$, Cm$^R$ | St 12 | This study |
| **St 237** | N315 ex w/o ϕ / pCNi-P$_{comGA}$-*comGA-mCh* / pRIT-P$_{comEA}$-*comEA-egfp* | Erm$^R$, Cm$^R$, Kan$^R$ | St228 | This study |
| **St 238** | N315 ex w/o ϕ / pCNi-P$_{comG}$-*comGA-mCh* / pRIT-P$_{comEC}$-*comEC-egfp* | Erm$^R$, Cm$^R$, Kan$^R$ | St228 | This study |
| **St 239** | N315 ex w/o ϕ / pCNi -P$_{comGA}$-*comGA-mCh* / pRIT-P$_{comFA}$-*comFA-egfp* | Erm$^R$, Cm$^R$, Kan$^R$ | St228 | This study |
| **St 240** | N315 ex w/o ϕ / pCNi -P$_{comGA}$-*comGA-mCh* / pRIT-P$_{dprA}$-*dprA-egfp* | Erm$^R$, Cm$^R$, Kan$^R$ | St228 | This study |
| **St 241** | N315 ex w/o ϕ / pCNi -P$_{comGA}$-*comGA-mCh* / pRIT-P$_{ssbC}$-*ssbB-egfp* | Erm$^R$, Cm$^R$, Kan$^R$ | St228 | This study |
| **St 242** | N315 ex w/o ϕ / pCNi -P$_{comGA}$-*comGA-mCh* / pRIT-P$_{comGA}$-*comGA-egfp* | Erm$^R$, Cm$^R$, Kan$^R$ | St228 | This study |
| **St 243** | N315 ex w/o ϕ / pRIT-P$_{comGA}$-*comGC-FLAG* | Erm$^R$, Cm$^R$ | St 12 | This study |
| **St 266** | N315 ex w/o ϕ / pCNi -P$_{comGA}$-*comGA-mCh* / pRIT-P$_{comGA}$-*comGC-FLAG* | Erm$^R$, Cm$^R$, Kan$^R$ | St 228 | This study |

| Plasmids | | |
|---|---|---|
| **Name** | **Description** | **Source** |
| **pRIT-P$_{comGA}$-*gfp*** | *E. coli – S. aureus* low-copy-number shuttle vector (*AmpR – CmR*). Expression of GFP under the control of comGA promoter (PcomGA). | (Morikawa *et al.*, 2012) |
| **pBCB-7-ChK** | Non-replicative plasmid (*AmpR – KanR).* Encoding mCherry gene flanked by C and N terminal linkers. | (Pereira *et al.*, 2010) |
| **pBCB-8-GK** | Non-replicative plasmid (*AmpR – KanR).* Encoding EGFP gene flanked by C and N terminal linkers. | (Pereira *et al.*, 2010) |
| **pCN34** | *E. coli – S. aureus* low-copy-number shuttle vector (*AmpR – KanR)* | (Charpentier *et al.*, 2004) |
| **pCN35** | *E. coli – S. aureus* high-copy-number shuttle vector (*AmpR – ErmR)* | (Charpentier *et al.*, 2004) |

#### Supplementary table 7. *S. aureus* late competence genes studied

| Gene | Locus tag | Gene length | Promoter | Promoter length |
|---|---|---|---|---|
| *comGA* | SA1374 | 975 bp | $P_{comGA}$ | 256 bp |
| *comGC* | SA1372 | 312 bp | $P_{comGA}$ | 256 bp |
| *comEA* | SA1418 | 687 bp | $P_{comEA}$ | 284 bp |
| *comEC* | SA1416 | 2202 bp | $P_{comEC}$ | 224 bp |
| *comFA* | SA0705 | 1083 bp | $P_{comFA}$ | 1815 bp |
| *dprA* | SA1092 | 873 bp | $P_{dprA}$ | 325 bp |
| *ssbC* | SA1899 | 396 bp | $P_{ssbC}$ | 193 bp |

#### Supplementary table 8. Primers used in this study

| Primer | Name | 5' oligonucleotide sequence | Amplified DNA | DNA template | Used to |
|---|---|---|---|---|---|
| P152 | KpnI-$P_{comGA}$-F | GCA GGT ACC GTT CGA TGA ATT CGC AGT TGT TGG AGA TAC | $P_{comGA}$-*comGA* | N315 genomic DNA | Create pBCB-$P_{comGA}$-*comGA-egfp* by ligating KpnI-digested $P_{comGA}$-*comGA* insert into KpnI-digested pBCB8-GK plasmid |
| P220 | KpnI-*comGA*-R | GCA GGT ACC AAT GTA TTT ATC CAT TGT AGT TTC ACA AAT GAC ACC TGC | | | |
| P320 | pBCB-*comGA*-F | GGA GGT GTT TTT TTG AAG ATT CTA TTT CAA GAA ATA ATT AAT AAA GCG | $P_{comGA}$-*comGA-egfp* | pBCB-$P_{comGA}$-*comGA-egfp* | Create pRIT-$P_{comGA}$-*comGA-egfp* by Gibson assembly |
| P321 | pRIT-*egfp*-SpeI-R | GCT ATG ACC ATG ATT ACG CCA AGC TGC ACT AGT GCT CAG CCG | | | |
| P322 | PstI-pRIT-F | CGG CTG AGC ACT AGT GCA GCT TGG CGT AAT CAT GGT CAT AGC | pRIT-$P_{comGA}$ | pRIT-$P_{comGA}$-*gfp* | |
| P323 | pRIT-$P_{comGA}$-R | CGC TTT ATT AAT TAT TTC TTG AAA TAG AAT CTT CAA AAA AAC ACC TCC | | | |
| P335 | pRIT-X-F | GGC TTA ACT ATG CGG CAT CAG AGC | Variable | All pRIT plasmids | Check insertions in pRIT plasmids |
| P336 | pRIT-X-R | GAA TTG TGA GCG GAT AAC AAT TTC ACA CAG G | | | |
| P376 | LinkerA-*egfp*-F | GGT ACC AGC GCT ATC GAT CGG CCG | pRIT-*egfp* | pRIT-$P_{comGA}$-*comGA-egfp* | Amplify pRIT-*egfp* linear plasmid fragment for Gibson assembly |
| P377 | EcoRI-pRIT-R | GAA TTC ACG AAC ACA TAT GGT GCA CTC TCA GTA CAA TCT GCT CTG ATG | | | |
| P378 | pRIT-EcoRI-$P_{comEA}$-F | ACC ATA TGT GTT CGT GAA TTC ATG GGA TGC TGT GCA AGG AGA TTT ACC | $P_{comEA}$-*comEA* | N315 genomic DNA | Create pRIT-$P_{comEA}$-*comEA-egfp* by Gibson assembly |
| P379 | LinkerA-*comEA*-R | CCG ATC GAT AGC GCT GGT ACC TAT CGT GAA ATA AGA TTT CAG TTT ATC AAA AGT TTT ACT | | | |
| P380 | pRIT-EcoRI-$P_{comFA}$1-F | ACC ATA TGT GTT CGT GAA TTC GGT GGA TTA GGT TTA GGC TAT GTT GGC | $P_{comFA}$1-*comFA* | N315 genomic DNA | Create pRIT-$P_{comFA}$1-*comFA-egfp* by Gibson assembly |
| P381 | Linker_A-*comFA*-R | CCG ATC GAT AGC GCT GGT ACC TTC ATC AAT CCA ACC TCT TTT TAA TGC TAA TTT GTT CAT | | | |
| P398 | LinkerA-*mCh*-F | CCG GAA GGA GAT ATA CAT ATG GCT ATC ATT AAA GAG TTC ATG CGC TTC AAA G | *mCh* | pBCB-7-ChK | Create pRIT-$P_{comGA}$-*comGA-mCh* by Gibson assembly |
| P399 | C-term-R | GCC TCG AGT GGT TGA GCA TGC | | | |
| P400 | C-term-F | AGG CCT GAT GCA TGC TCA ACC | pRIT-$P_{comGA}$-*comGA* | pRIT-$P_{comGA}$-*comGA-egfp* | |
| P401 | *mCh*-LinkerA-R | GCA TGA ACT CTT TAA TGA TAG CCA TAT GTA TAT CTC CTT CCG GCC GAT CGA TAG | | | |
| P454 | pRIT-EcoRI-$P_{comEC}$-F | TGC ACC ATA TGT GTT CGT GAA TTC ACA AGG TGT ATC TAC TGA AGG TGC | $P_{comEC}$-*comEC* | N315 genomic DNA | Create pRIT-$P_{comEC}$-*comEC-egfp* by Gibson assembly |
| P455 | LinkerA-*comEC*-R | CCG ATC GAT AGC GCT GGT ACC TAA ACC ACT TGC ATT TCC ATA AGA GTT TG | | | |
| P456 | pRIT-EcoRI-$P_{dprA}$-F | TGC ACC ATA TGT GTT CGT GAA TTC AAT AAC AGG CGT CTT TAG TAA TGA GGC | $P_{dprA}$-*dprA* | N315 genomic DNA | Create pRIT- $P_{dprA}$-*dprA-egfp* by Gibson assembly |
| P457 | LinkerA-*dprA*-R | CCG ATC GAT AGC GCT GGT ACC AAT ATA GTA GTC TTC AAA TAT ATC ATT AGC GTT TAA | | | |
| P460 | pRIT-EcoRI-$P_{ssbC}$-F | C ACC ATA TGT GTT CGT GAA TTC ATT CCC ACA TCC CAT TAG TTT AAT ATT TAT GAT TTT TG | $P_{ssbC}$-*ssbC* | N315 genomic DNA | Create pRIT-$P_{ssbC}$-*ssbC-egfp* by Gibson assembly |

| | | | | | |
|---|---|---|---|---|---|
| P461 | LinkerA-*ssbC*-R | CCG ATC GAT AGC GCT GGT ACC AAT TTC TAA TAA GTC ATG ATT ATC TAT ATT TTG AGA | | | |
| P505 | *aphA3*-F | GTT TAA GGG CCC ACC TAG GGG TTT CAA AAT CGG CTC CG | *aphA3* | pCNi | Check for kanamycin resistance cassette |
| P506 | *aphA3*-R | CTA TGA CTC GAG GCC GCG GCG CTC GGG ACC CCT ATC TAG C | | | |
| P519 | *comEA*-pRIT-F | CTG AAA TCT TAT TTC ACG ATA TAA GCA CTA GTG CAG CTT GGC GTA ATC | pRIT-$P_{comEA}$ | pRIT-$P_{comEA}$-*comEA-egfp* | Create pRIT-$P_{comEA}$-*egfp-comEA* by Gibson assembly |
| P520 | *egfp*-$P_{comEA}$-R | GCT AGC CAT TCC TCA ACT TTA TAC ACG TCT GAG CGA GTG | | | |
| P521 | $P_{comEA}$-*egfp*-F | CTC AGA CGT GTA TAA AGT TGA GGA ATG GCT AGC AAA GGA GAA GAA CTT TTC ACT GGA | *egfp* | pBCB8-GK | |
| P522 | *comEA-egfp*-R | TAA AAA TTG ATA CAA TAA AAC CAC GCC TCG AGT GGT TGA GCA TGC ATC | | | |
| P523 | *egfp-comEA*-F | CCA CTC GAG GCG GTG GTT TTA TTG TAT CAA TTT TTA TTA CGC TAT AAA GAT TTT TTA ACT | *comEA* | N315 genomic DNA | |
| P524 | pRIT-*comEA*-R | AAG CTG CAC TAG TGC TTA TAT CGT GAA ATA AGA TTT CAG TTT ATC AAA AGT TTT ACT TCC | | | |
| P525 | pRIT-$P_{comFA}$2-F | ATA TGT GTT CGT GAA TTC CAG TTT GAT CAT AAT TCA GTG TTA CTA TAC ATG GTA CTG | $P_{comFA}$2 | N315 genomic DNA | Create pRIT-$P_{comFA}$2-*comFA-egfp* by Gibson assembly |
| P526 | *comFA*-$P_{comFA}$2-R | CGA ACT CTC TGT TAT TTT ATA TCT TGT TAC ATT ATC CAT TCG AC | | | |
| P527 | *comFA*-F | ATG GAT AAT GTA ACA AGA TAT AAA ATA ACA GAG AGT TCG CAA AGT TCA TCA CAA GC | pRIT-*comFA-egfp* | pRIT-$P_{comFA}$1-*comFA-egfp* | |
| P528 | $P_{comFA}$2-pRIT-R | TGA ATT ATG ATC AAA CTG GAA TTC ACG AAC ACA TAT GGT GCA CTC TCA GTA CAA TCT G | | | |
| P531 | SalI-$P_{comGA}$-F | TGT GTC GAC TGT TCG TGA ATT CGC AGT TGT TGG AGA TAC ATT ATT TAA TAA TGG | $P_{comGA}$-*comGA-mCh* | pRIT-$P_{comGA}$-*comGA-mCh* | Create pCNi-$P_{comGA}$-*comGA-mCh* by ligating SalI-SacI-digested $P_{comGA}$-*comGA-mCh* insert into SalI-SacI-digested pCNi plasmid |
| P532 | SacI-*mCh*-R | TTA GAG CTC TCA GCC GGC GCG GC | | | |
| P533 | pCN-X-F | TGG TAT CTT TAT AGT CCT GTC GGG TTT CGC | Variable | All pCN plasmids | Check insertions in pCN plasmids |
| P534 | pCN-X-R | ATT TAG TTT TGG TTC ATC TTC TGT TAA CTT ATT AAC TCT TTC CGC | | | |
| P560 | *comGC*-FLAG-pRIT-F | GCA AAT GAC TAC AAG GAC GAC GAT GAC AAG TGA GCA CTA GTG CAG CTT GGC | pRIT-$P_{comGA}$ | pRIT-$P_{comGA}$-*comGA-egfp* | Create pRIT-$P_{comGA}$-*ComGC*-FLAG by Gibson assembly |
| P561 | *comGC*-$P_{comGA}$-R | TTG AGT TTT CTT AAG AAA TTT AAA CAT AAA AAC ACC TCC TAC ATA TAA TCA CGT AGG AGG | | | |
| P562 | $P_{comGA}$-*comGC*-F | GTA GGA GGT GTT TTT ATG TTT AAA TTT CTT AAG AAA ACT CAA GCG TTT ACA TTG ATA GAG | *ComGC*-FLAG | N315 genomic DNA | |
| P567 | FLAG-*comGC*-R | GTC ATC GTC GTC CTT GTA GTC ATT TGC AAC TGC TTC TCC ATT ACT AAT TGT TAT TGT CTC | | | |

### Supplementary videos

**Supplementary video 1. 360 ° rotation of a *S. aureus* competent cell with cytoplasmic ComGA-EGFP.**

Strain St113 (pRIT-$P_{comGA}$-*comGA-egfp*) was grown for 25h ($10^{-5}$ dilution) in CS2 medium. ComGA appears in the cytoplasm.

**Supplementary video 2. 360 ° rotation of a *S. aureus* competent cell with ComGA-EGFP associated to the inner face of the membrane.**

Strain St113 (pRIT-$P_{comGA}$-*comGA-egfp*) was grown for 22h ($10^{-5}$ dilution) in CS2 medium. ComGA appears associated to the inner face of the membrane.

**Supplementary video 3. 360 ° rotation of a *S. aureus* competent cell with a ComGA-EGFP focus.**

Strain St113 (pRIT-$P_{comGA}$-*comGA-egfp*) was grown for 22h ($10^{-5}$ dilution) in CS2 medium. ComGA appears as a focus.

**Supplementary video 4. 360 ° rotation of a *S. aureus* competent cell harboring a ComGA-mCherry focus and stained with Vanco-BODIPY.**

Strain St228 (pRIT-$P_{comGA}$-*comGA-mCh*) was grown for 22h ($10^{-5}$ dilution) in CS2 medium and stained with Vancomycin BODIPY FL (Vanco-BODIPY). The cell wall is stained by the Vanco-BODIPY in green and a ComGA-mCherry focus appears in red.
